# Trajectory Uncertainty Framework (TUF): A Modular Framework for Identifying Transitional and Branch-Point Cell States in Single-Cell Trajectory Analysis

**DOI:** 10.64898/2026.07.31.742012

**Authors:** Masoud Mahdavifar, Zahra Mohammadifar, Tara Iranpourtari

## Abstract

Single-cell trajectory inference methods assign pseudotime coordinates but provide limited information on assignment uncertainty, especially at transitional states and branch points. We introduce the Trajectory Uncertainty Framework (TUF), a modular, downstream approach that defines a 2D uncertainty coordinate system for single-cell data. TUF decomposes local uncertainty into the Temporal Entropy Score (TES; temporal heterogeneity) and the Trajectory Divergence Score (TDS; directional divergence).

Synthetic benchmarks demonstrate that the joint interpretation of TES and TDS helps distinguish true fate bifurcations. Applied to pancreatic endocrinogenesis, intestinal epithelium, glioblastoma, and breast cancer datasets, the TUF coordinate system identifies known transitional populations. To isolate the transcriptional drivers of uncertainty beyond baseline tumor biology, we employed a fractional logit residual analysis. This reveals that TES and TDS are associated with context-specific transitional programs: in glioblastoma, residual TES is enriched for inflammatory remodeling while TDS marks proliferative and metabolic stress, whereas in breast cancer, residual TES is enriched for EMT-associated and extracellular matrix remodeling programs, and TDS marks distinct lineage-associated programs. These axes show minimal gene overlap (Jaccard 0.079 in glioblastoma; 0.028 in breast cancer) and remain robust across independent Julia and Python implementations.

SiCell.jl provides an efficient, open-source implementation of TUF and is available under the MIT license via the Julia General Registry and GitHub.

## 1 Introduction

### 1.1 Trajectory inference in single-cell biology

Single-cell RNA sequencing (scRNA-seq) enables the reconstruction of dynamic cellular processes from snapshot data, typically through trajectory inference methods that organize cells along continuous developmental paths Jovic et al. (2022). A wide range of computational approaches have been proposed for this purpose, including graph-based methods, diffusion-based models, and principal curve or tree-based approaches Zhang et al. (2023). While these methods successfully recover coarse-grained differentiation topology, accurately resolving fine-grained transitional states and ambiguous branching regions remains challenging. Biologically, these regions may represent intermediate cell states, where neighboring cells temporarily share transcriptional features while still following the same developmental path Saelens et al. (2019).

### 1.2 Limitations of pseudotime-based representations

Most trajectory inference methods rely on pseudotime, which assigns each cell a scalar value representing its position along an inferred developmental progression Saelens et al. (2019). While effective for ordering cells globally, pseudotime representations inherently provide only deterministic mappings from high-dimensional transcriptomic space to a one-dimensional continuum. Consequently, these methods often fail to capture local ambiguity in developmental decisions, particularly in regions of trajectory bifurcation where multiple lineage fates are simultaneously plausible. This limitation is amplified by noise, sparsity, and sampling bias in scRNA-seq data.

### 1.3 A 2D coordinate system for quantifying uncertainty

Existing approaches for uncertainty estimation remain limited, often relying on heuristic thresholds, distance-based criteria, or single metrics that cannot distinguish different sources of ambiguity Saelens et al. (2019). To address this, we introduce the Trajectory Uncertainty Framework (TUF), a modular approach that operates downstream of existing trajectory inference workflows. Rather than proposing a single uncertainty score, TUF defines a 2D uncertainty coordinate system using two simple, interpretable local metrics: the Temporal Entropy Score (TES), which quantifies local temporal heterogeneity, and the Trajectory Divergence Score (TDS), which quantifies local directional divergence. Individually, these metrics capture distinct geometric features; jointly, they provide a low-redundancy coordinate system for interpreting trajectory uncertainty.

### 1.4 Software implementations and accessibility

To facilitate adoption, we provide open-source software implementations. The framework is available as the Julia package SiCell.jl, as well as reproducible Python scripts with native support for AnnData objects.

## 2 Methods

### 2.1 Standard trajectory inference workflow

TUF is designed to operate downstream of conventional single-cell trajectory inference workflows. Raw count matrices are first processed using standard preprocessing procedures. A k-nearest neighbor (kNN) graph is subsequently constructed and used for trajectory inference and pseudotime estimation. Because TUF is agnostic to the upstream method, it can be applied to outputs generated by a variety of approaches, including Diffusion Pseudotime (DPT) Haghverdi et al. (2016). The resulting pseudotime values, neighborhood graph structure, and diffusion embedding serve as inputs to TUF (Figure 1).

**Figure 1:**
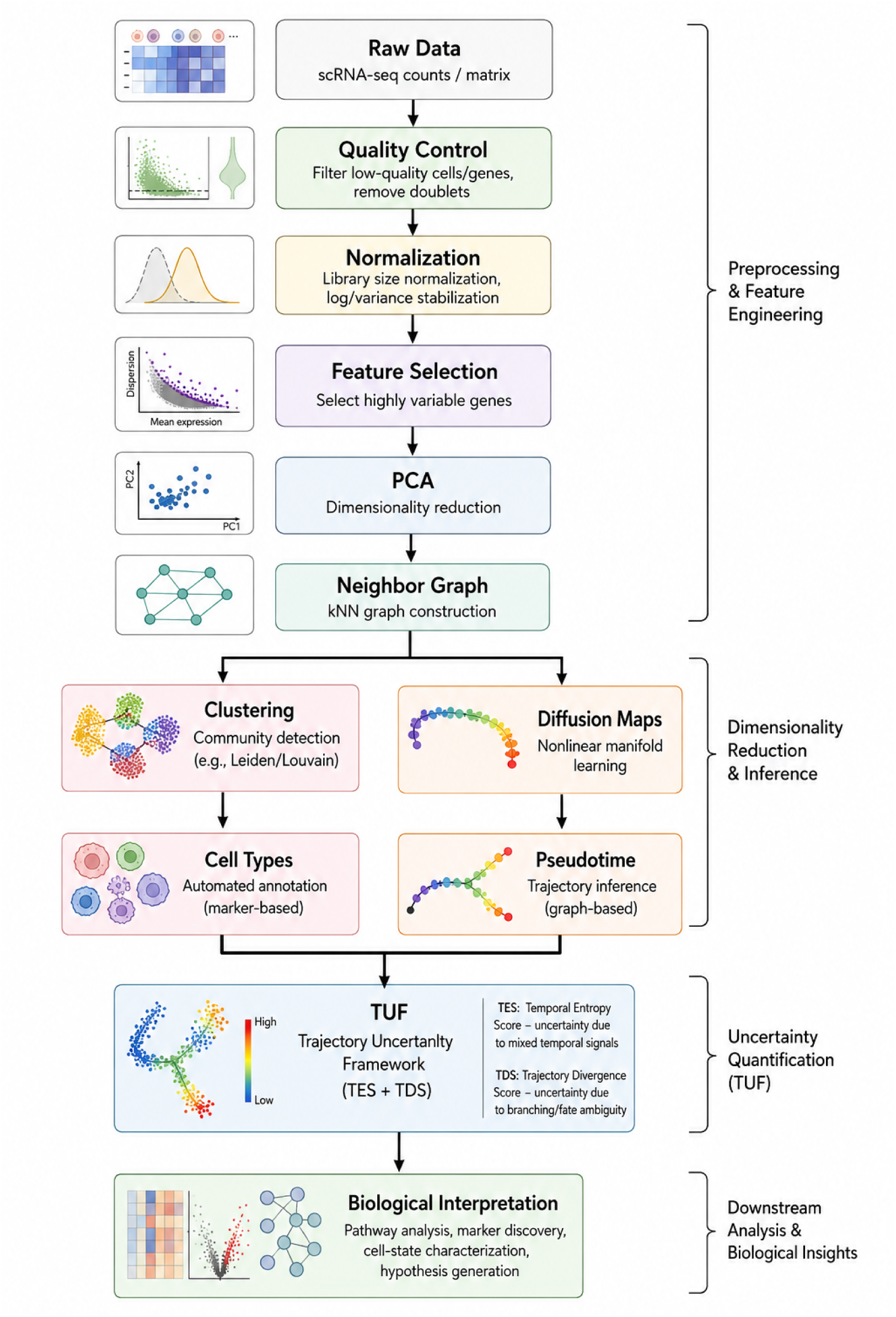
Standard single-cell analysis workflow feeding into TUF.

### 2.2 The Trajectory Uncertainty Framework (TUF)

TUF computes two complementary per-cell metrics directly from the k-nearest neighbor graph and diffusion embedding. TES requires sorting the local pseudotime values and therefore scales as *O*(*Nk* log *k*). TDS is computed in a single pass over the neighborhood and scales as *O*(*Nk*). Consequently, the overall complexity of TUF is dominated by TES and is *O*(*Nk* log *k*).

#### 2.2.1 Temporal Entropy Score (TES)

For each cell *i*, TES quantifies local temporal heterogeneity by measuring pairwise pseudotime variability among its *k* nearest neighbors *N* (*i*):

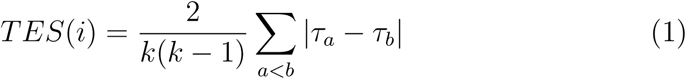

where *τ_a_* and *τ_b_* denote pseudotime values of neighboring cells. **Crucially, pseudotime must be min-max scaled to the** [0, 1] **interval prior to TUF application to ensure TES is bounded in** [0, 1]. To improve computational efficiency, TES can be evaluated in *O*(*k* log *k*) time per cell using the identity for the Gini mean difference:

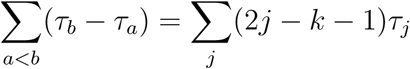

over sorted neighbor pseudotime values. Mathematically, TES corresponds to the normalized average pairwise pseudotime difference (equivalent to a scaled Gini mean difference). We use this formulation because it provides a continuous, parameter-free measure of local temporal heterogeneity without requiring discretization into histogram bins, while the term “temporal entropy” reflects the conceptual interpretation of increasing pseudotemporal uncertainty arising from heterogeneous neighborhood pseudotime.

#### 2.2.2 Trajectory Divergence Score (TDS)

TDS quantifies directional dispersion of local neighborhood structure in embedding space. For each cell *i*, unit displacement vectors are computed from *i* to each neighbor *j ∈ N* (*i*):

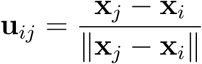

The mean resultant vector length measures directional coherence:

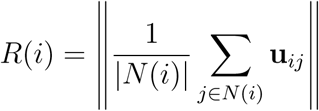

TDS is then defined as:

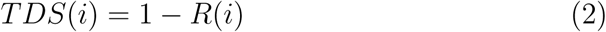

TDS approaches 0 when neighbors are aligned in a consistent direction (committed lineages), and approaches 1 when neighbor directions are maximally dispersed. Because TDS relies on angular relationships, it requires an embedding where local distances and angles are geometrically meaningful (e.g., diffusion components or PCA space). **We explicitly caution that nonlinear visualization coordinates such as UMAP or t-SNE are unsuitable inputs for TDS calculation, as their non-metric distortions produce meaningless directional variances.** Additionally, TDS is sensitive to boundary artifacts where small displacement magnitudes produce large angular variance after normalization; therefore, we recommend applying a small magnitude threshold *ɛ* to filter out near-zero displacement vectors, or relying on quantile-based thresholding for the final TDS scores. This formulation is closely related to directional statistics Jammalamadaka and Sengupta (2001); Mardia and Jupp (2009).

#### 2.2.3 The TES/TDS coordinate system

TES measures local temporal heterogeneity and is sensitive to regions where neighboring cells occupy a broad range of developmental stages. TDS quantifies directional divergence and is sensitive to local angular incoherence. While high TDS occurs at fate bifurcations, it is also triggered by spatial crossings or convergence funnels. Therefore, TDS alone is not a specific marker of branching; rather, it is a directional incoherence metric. The core contribution of TUF is the joint interpretation of these metrics as a 2D coordinate system (Figure 2D). True bifurcations are characterized by high TDS combined with low TES, whereas spatial crossings show high TDS combined with elevated TES. Conversely, regions exhibiting high TES but low TDS represent temporal heterogeneity without directional divergence (e.g., locally asynchronous differentiation along a shared lineage path), while low scores on both metrics denote committed, coherent lineages.

**Figure 2:**
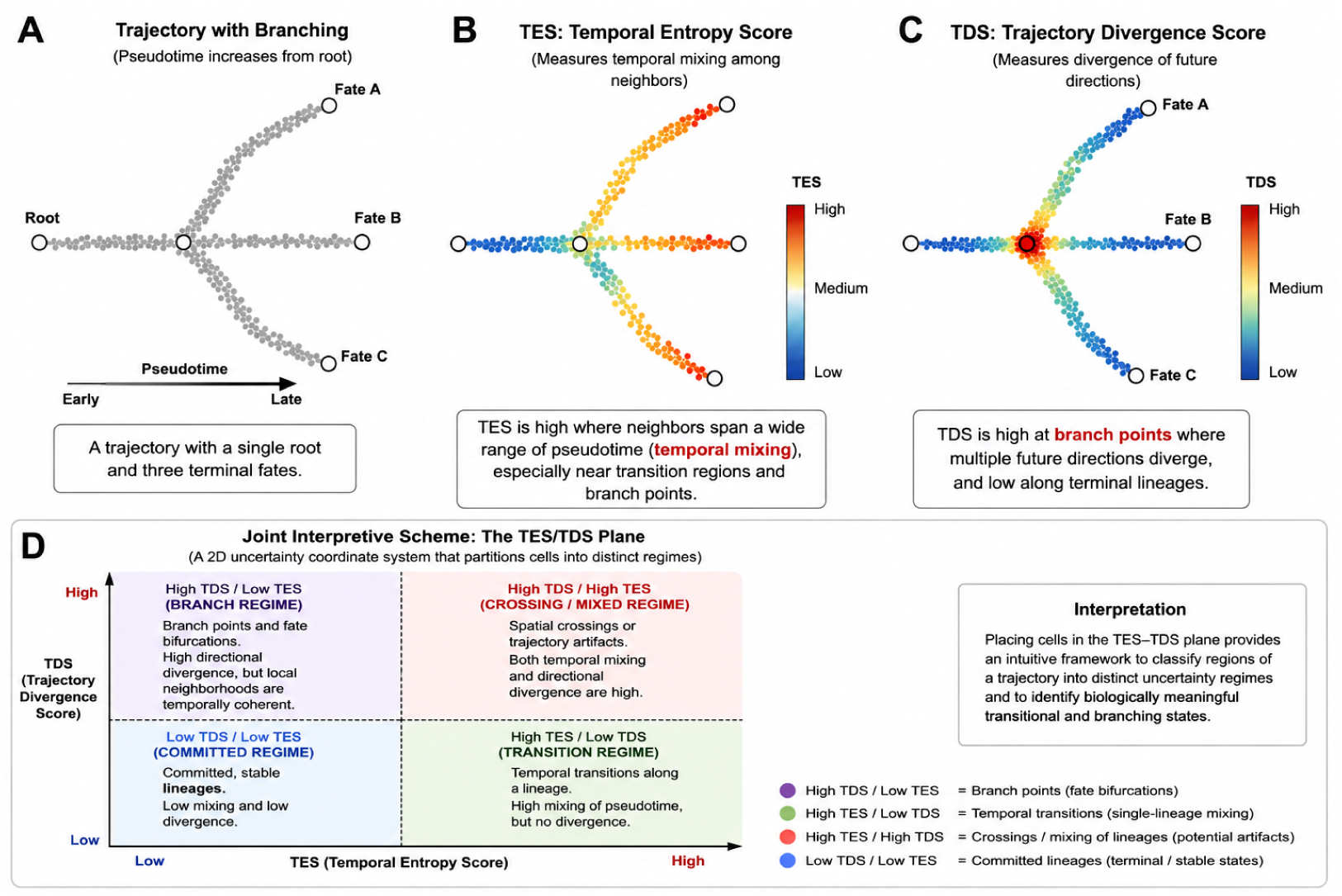
The 2D uncertainty coordinate system defined by TUF. (A) Example developmental trajectory. (B) TES captures temporal heterogeneity. (C) TDS captures directional divergence. (D) Joint interpretive scheme: The TES/TDS plane partitions cells into distinct uncertainty regimes. High TES/low TDS are consistent with temporal transitions; High TDS/low TES are consistent with fate bifurcations; High scores on both denote spatial crossings/artifacts; Low scores on both denote committed lineages.

### 2.3 Identification of transition-associated programs

To characterize molecular signatures associated with trajectory uncertainty beyond known biological confounders, we employed a residual-based approach. We calculated signature scores for hypoxia, stemness, cell cycle, and epithelial-mesenchymal transition (EMT) based on canonical gene sets using standard mean-expression aggregation, which were subsequently minmax scaled to the [0, 1] interval to match the dependent variables. These four signatures were chosen as they represent the most prevalent pan-cancer confounders known to covary with differentiation states.

Because TES and TDS are bounded in [0, 1], we employed a fractional logit regression framework Papke and Wooldridge (1996) to partition variance. Model fit was evaluated using the squared Pearson correlation between observed and predicted values. The unexplained residual for each cell was then calculated on the raw scale as (Residual = Observed − Predicted).

We defined the “high residual” group as the top 5% of cells exhibiting the largest positive residuals across the dataset, restricting selection to cells whose observed uncertainty substantially exceeds confounder predictions. Differential expression was performed comparing this group against the background using a Wilcoxon rank-sum test (FDR *<* 0.05, log_2_FC *>* 0.25). Over-representation analysis (ORA) was performed using Enrichr Kuleshov et al. (2016) against the GO Biological Process Ashburner et al. (2000), MSigDB Hallmark Liberzon et al. (2015), and Reactome libraries. ORA was chosen over GSEA because it is specifically suited for highly filtered gene lists derived from extreme-tail subset comparisons, whereas GSEA evaluates the entire ranked distribution which dilutes the subset signal. Low redundancy between TES and TDS gene sets was assessed using the Jaccard index against 10,000 size-matched random gene sets.

### 2.4 Datasets

Datasets are publicly available from their original publications (Table 1). For the primary biological analyses, the neighborhood size was set to *k* = 20 and TDS was computed using the top 10 diffusion components.

**Table 1:**
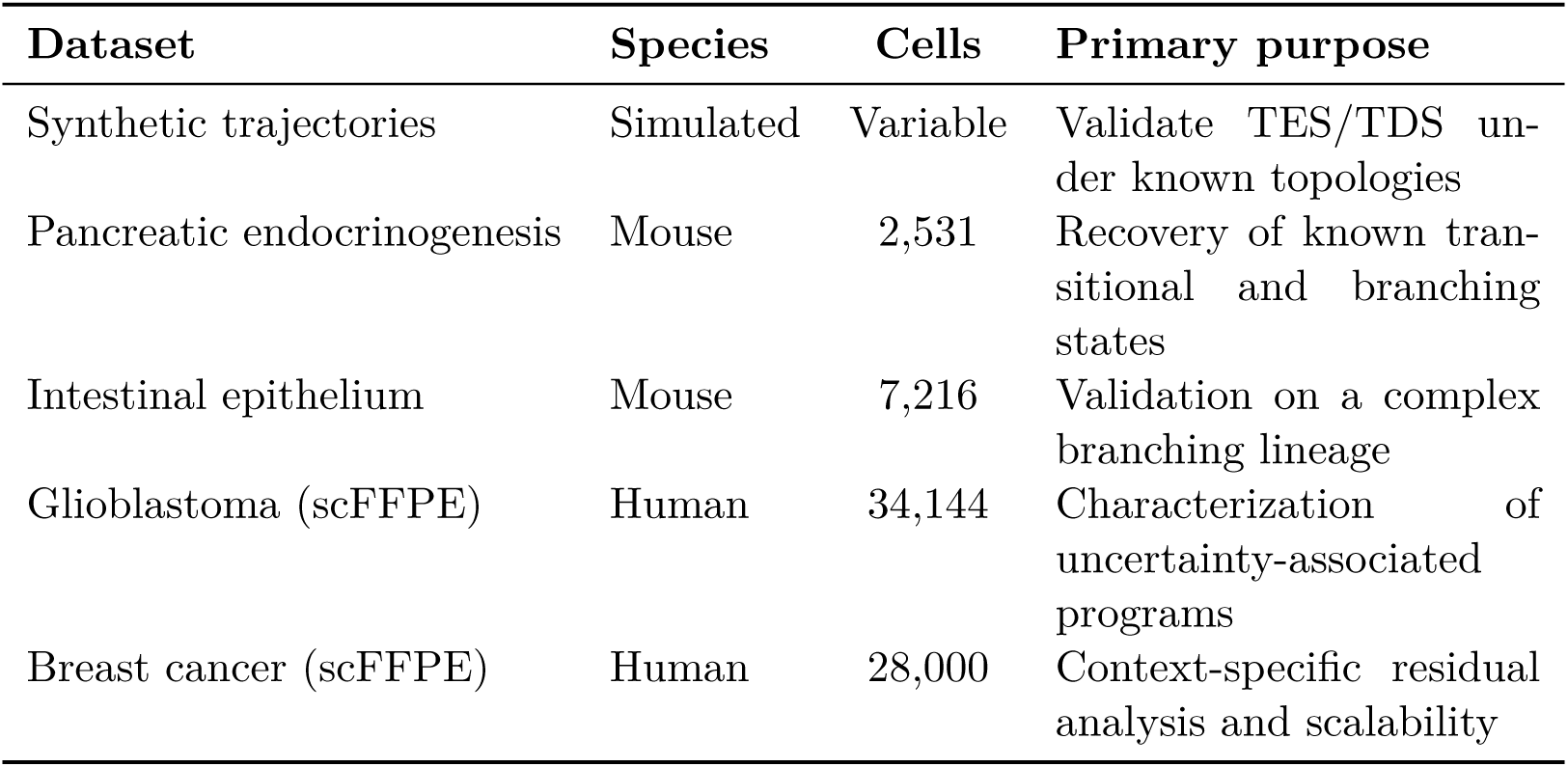
Datasets used throughout this study and their role in evaluating TUF.

| Dataset | Species | Cells | Primary purpose |
| --- | --- | --- | --- |
| Synthetic trajectories | Simulated | Variable | Validate TES/TDS under known topologies |
| Pancreatic endocrinogenesis | Mouse | 2,531 | Recovery of known transitional and branching states |
| Intestinal epithelium | Mouse | 7,216 | Validation on a complex branching lineage |
| Glioblastoma (scFFPE) | Human | 34,144 | Characterization of uncertainty-associated programs |
| Breast cancer (scFFPE) | Human | 28,000 | Context-specific residual analysis and scalability |

## 3 Results

### 3.1 TES isolates temporal mixing, while TDS alone conflates branch points with spatial crossings

To establish the baseline behavior of TES and TDS, we applied TUF to synthetic trajectories with known ground-truth topologies: a linear trajectory (negative control), crossing trajectories, a wide-angle Y-branch, and an asynchronous convergence. Each topology was evaluated across five independent random seeds. Summary statistics are reported in Table 2.

**Table 2:** Summary statistics for TES and TDS across synthetic ground-truth topologies (mean *±* std over 5 random seeds).

| Topology | Mean TES | Max TES | Mean TDS | Max TDS |
| --- | --- | --- | --- | --- |
| Linear (negative control) | $0.007 \pm 0.000$ | $0.010 \pm 0.000$ | $0.012 \pm 0.000$ | $0.215 \pm 0.070$ |
| Crossing trajectories | $0.031 \pm 0.002$ | $0.304 \pm 0.004$ | $0.502 \pm 0.001$ | $0.993 \pm 0.005$ |
| Y-branch (true bifurcation) | $0.012 \pm 0.000$ | $0.017 \pm 0.001$ | $0.163 \pm 0.004$ | $0.890 \pm 0.027$ |
| Asynchronous convergence | $0.016 \pm 0.001$ | $0.269 \pm 0.001$ | $0.173 \pm 0.003$ | $0.831 \pm 0.054$ |

TES remained near zero along the true Y-branch (mean = 0.012), as expected for cells sharing similar pseudotime, but rose sharply at spatial crossings (max = 0.304). In contrast, TDS responded strongly to both genuine branch points and spatial crossings/convergences. While the Y-branch produced high maximum TDS (0.890), crossing trajectories reached even higher values (0.993). This demonstrates that *TDS alone is not a specific marker of lineage branching*; it detects local directional incoherence regardless of its cause.

Notably, the linear negative control showed a surprisingly high max TDS (0.215 *±* 0.070), driven by boundary artifacts where small displacements produce large angular variance after normalization. This further motivates the use of magnitude filtering during TDS computation or robust, quantile-based thresholding of the final scores.

Critically, the joint TES/TDS coordinate system resolves these ambiguities: true bifurcations are characterized by high TDS + low TES, whereas geometric or embedding-induced crossings produce high values in both metrics. These synthetic topologies serve as geometric stress tests to isolate algorithmic behavior, rather than models of biological reality.

Figure 3 To further validate branching behavior in the absence of crossing artifacts, we tested a multi-fate star topology and a pseudotime noise sweep (Figure 4 and Supplementary Figure S1). These experiments confirmed that TDS scales with the number of diverging lineages at branch points, while TES remains largely insensitive to branching but sensitive to pseudotime noise.

**Figure 3:**
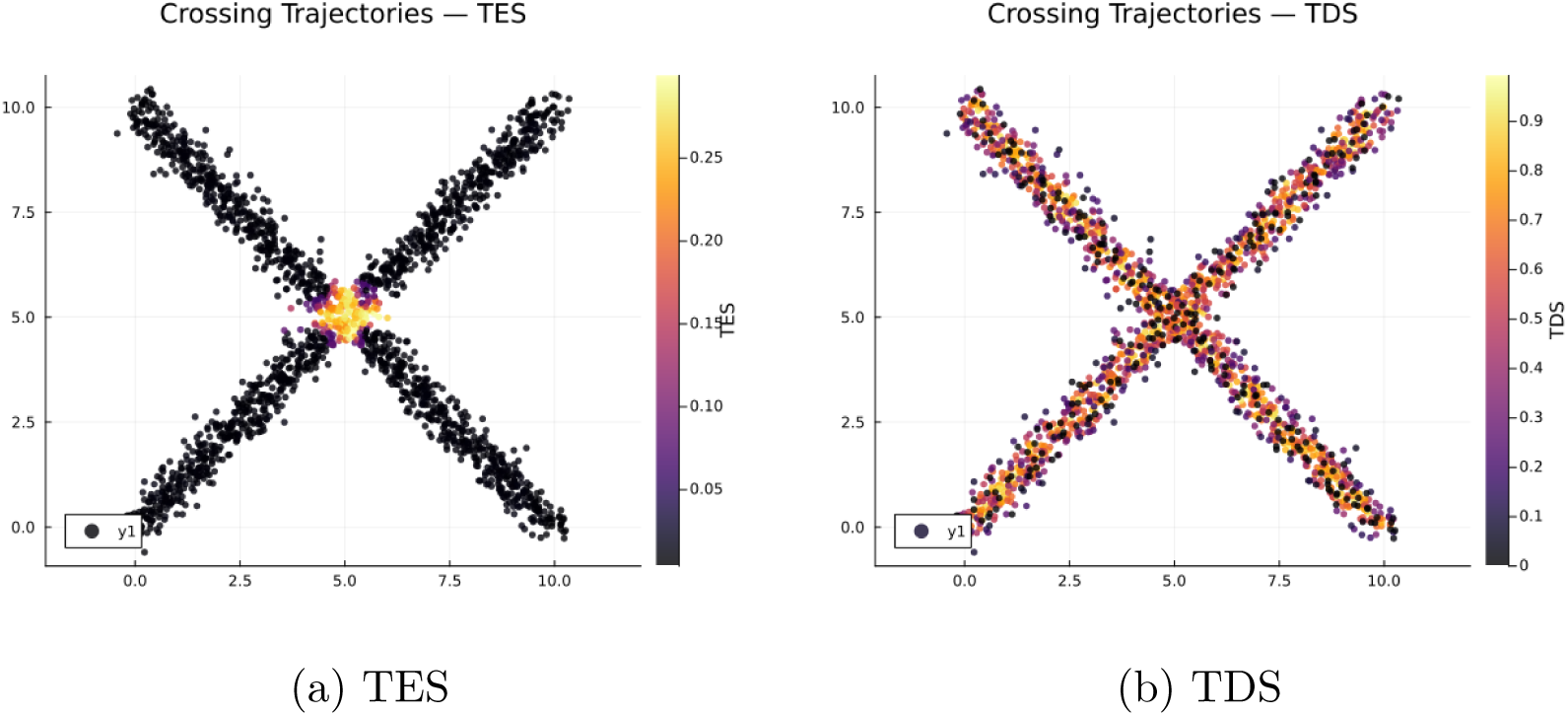
Spatial localization of trajectory uncertainty metrics on the crossing-trajectories synthetic dataset. (A) TES is sharply confined to the intersection region. (B) TDS is elevated at the intersection, reflecting spatial directional incoherence introduced by crossing lineages rather than true fate bifurcation.

**Figure 4:**
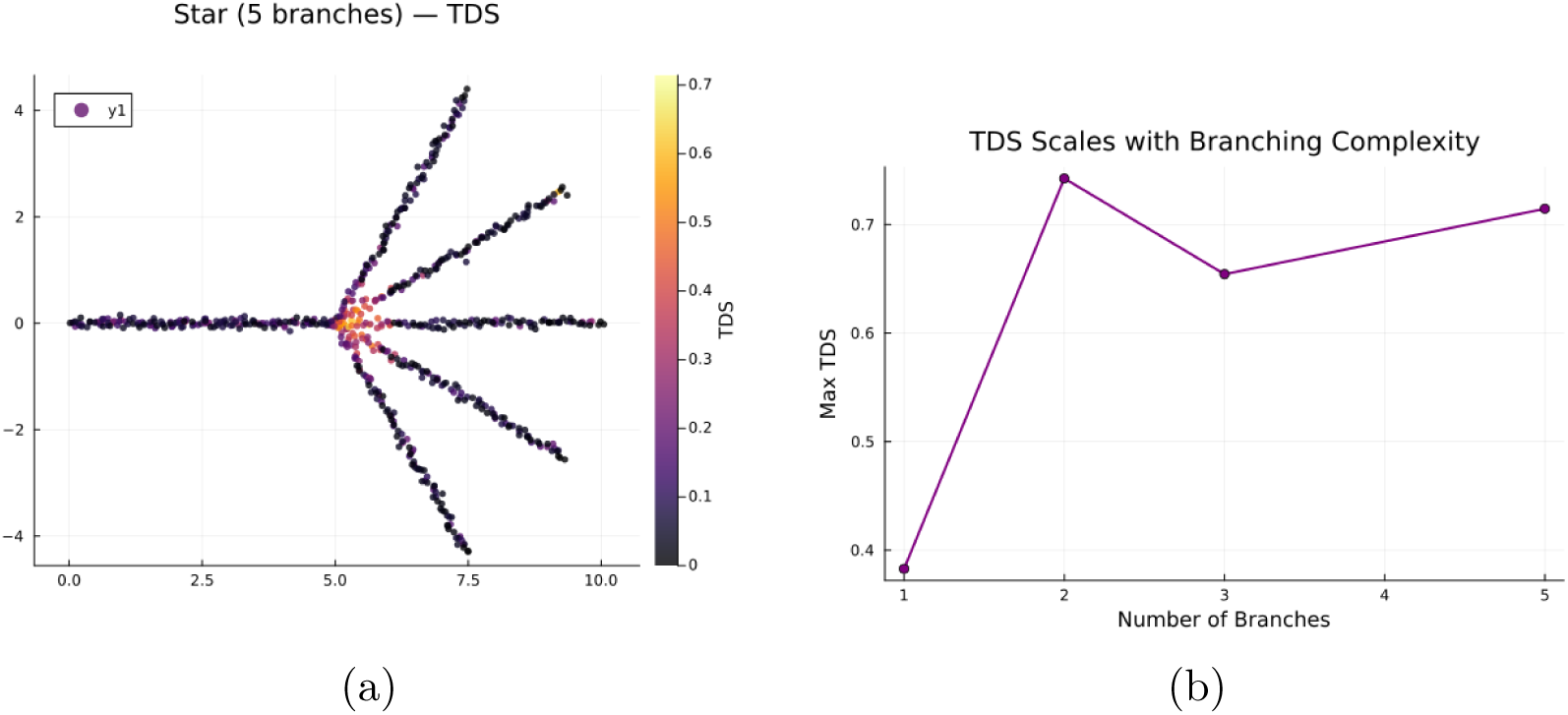
TDS as a branch-point detector in the absence of spatial-crossing confounds. (A) In a five-fate star topology, TDS localizes sharply to the branch hub. (B) Mean TDS at the hub increases with the number of branches.

Taken together, these results show that TES and TDS provide complementary information. Their joint use enables more accurate and specific identification of transitional and branch-point cell states.

### 3.2 TES captures continuous variance missed by discrete baselines

A natural question is whether TES provides information beyond simpler heuristic metrics, such as local neighborhood entropy. We computed neighborhood entropy as the Shannon entropy of discrete cluster labels within the kNN graph. Unlike TES, which operates directly on continuous pseudotime values, neighborhood entropy relies on discrete clustering assignments. Consequently, entropy is highly sensitive to clustering resolution and fails to capture intra-cluster temporal transitions.

To quantify this distinction, we correlated TES against these baselines in the pancreatic endocrinogenesis and glioblastoma datasets. TES exhibited weak positive correlation with neighborhood entropy (Spearman *ρ* = 0.121 and 0.112, respectively) and weak negative correlation with raw pseudotime (*ρ* = −0.158 and −0.346). TDS similarly showed weak negative correlation with pseudotime (*ρ* = −0.384 and −0.399). The low correlations—particularly between TES and neighborhood entropy—are expected. They reflect the fact that continuous temporal heterogeneity is largely distinct from discrete label mixing. TES bypasses the dependency on arbitrary clustering resolutions and is able to highlight continuous transitional dynamics that discrete baselines inherently miss.

### 3.3 TUF exhibits cross-pipeline consistency and computational robustness

To validate computational portability, we replicated the top 5% differential expression pipeline in an independent Python/Scanpy stack. The high-TES gene sets identified in Python were nearly identical to those in Julia. Furthermore, the low overlap between the TES and TDS gene sets (Jaccard *≈* 7.9%) was perfectly preserved across both languages, confirming that the complementary nature of the metrics is an intrinsic property of the algorithms. TUF was also robust to QC covariates (all absolute Spearman correlations *<* 0.04; Supplementary Table S1).

### 3.4 TES and TDS are associated with context-specific biological programs

We applied TUF to glioblastoma 10x Genomics (2025), breast cancer 10x Genomics (2024), pancreatic endocrinogenesis Bastidas-Ponce et al. (2019), and intestinal epithelium Haber et al. (2017) datasets. In pancreatic endocrinogenesis, high-TDS cells localized to the early Ngn3-low endocrine progenitor state, while high-TES cells mapped to late endocrine populations (including Epsilon, Fev+, and Delta cells) (Figure 5). In intestinal epithelium, high-TDS cells localized to progenitor compartments associated with lineage divergence van der Flier and Clevers (2009), whereas high-TES cells highlighted regions of local pseudotemporal heterogeneity, including secretory-associated populations such as Paneth and goblet cells (Figure 6).

**Figure 5:**
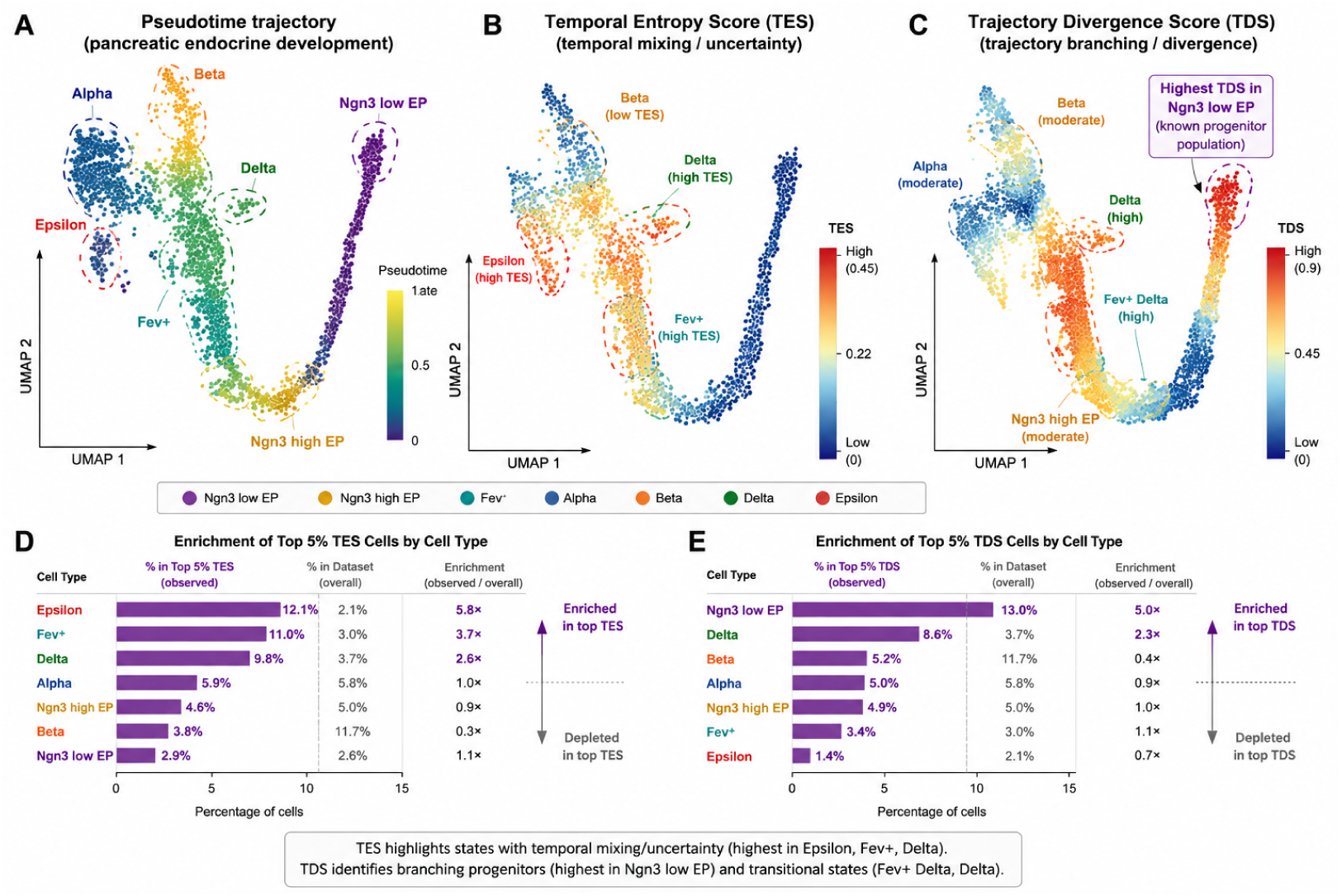
TUF recovers known transitional and branch-point populations in pancreatic endocrinogenesis. (A) Pseudotime trajectory on UMAP. (B) TES highlights regions of high temporal mixing. (C) TDS identifies branching progenitors. (D–E) Cell-type enrichment of top 5% TES and TDS cells.

**Figure 6:**
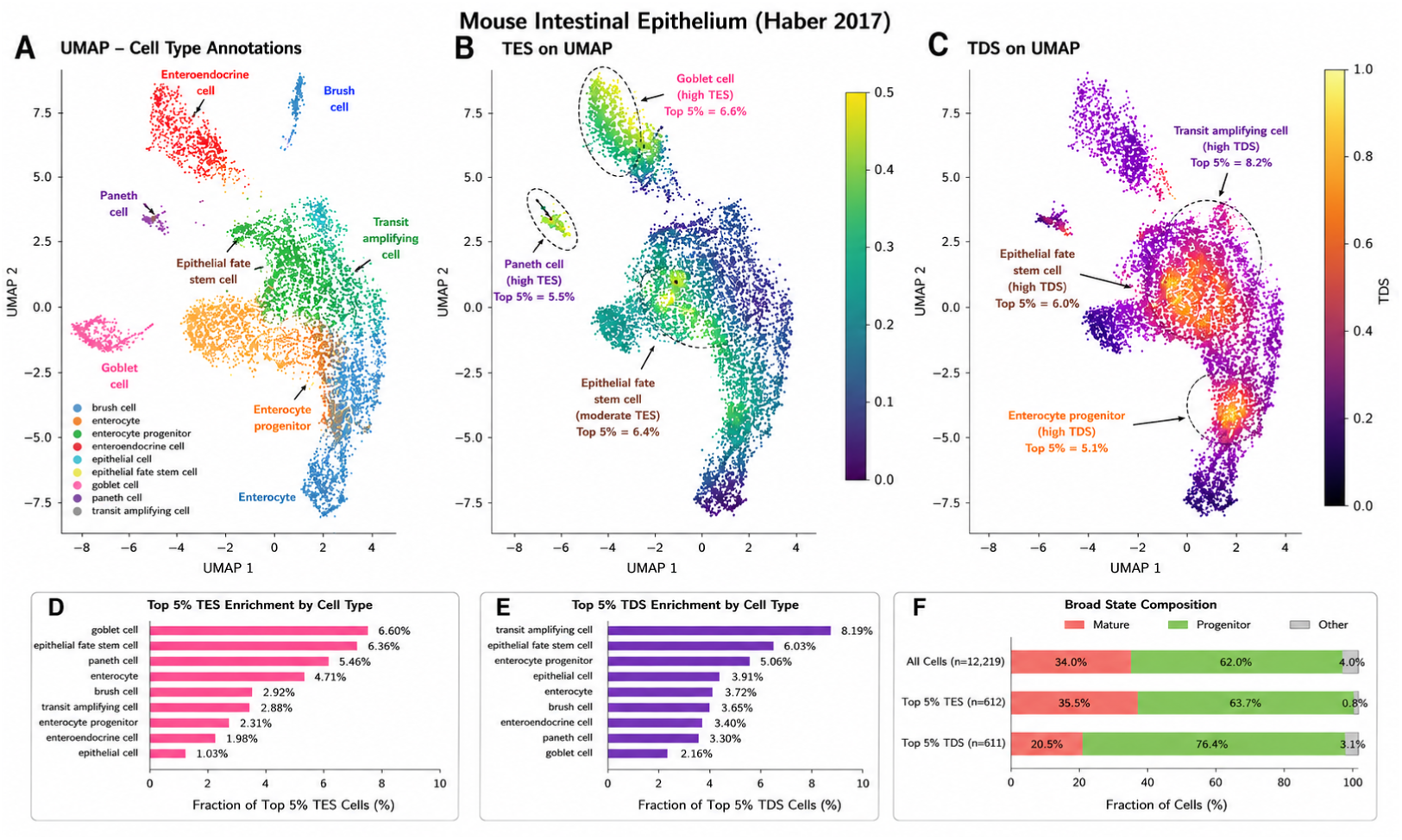
TUF applied to mouse intestinal epithelium (Haber et al., 2017). (A–C) UMAP annotations and TES/TDS localization. (D–F) Enrichment and progenitor composition of high-uncertainty cells.

Standard analysis of top 5% high-TES cells in glioblastoma yielded 343 genes enriched for broad stress pathways (Hypoxia, TNF-*α* signaling). High-TDS cells yielded 63 genes enriched for proliferative and metabolic programs. These gene sets showed low redundancy (Jaccard 0.079; permutation test mean null Jaccard = 0.154, SD = 0.009; *p <* 1 *×* 10*^−^*^4^).

We then applied fractional logit residual analysis. In GBM, residual TES cells were enriched for innate immune and inflammatory signaling (TNF-via NF-*κ*B, TGF-*β*, IL17; FDR *<* 10*^−^*^20^), alongside markers of phenotypic remodeling and the loss of mature structural glial programs (depletion of cilium assembly and axoneme genes Ho et al. (2009)). Top marker genes included the JAK/STAT feedback regulator *SOCS3* and immune modulator *AHR*. This profile—simultaneous inflammatory activation and loss of terminal identity—is consistent with cells actively undergoing temporal state transitions.

In contrast, residual TDS cells were enriched for a remarkably coherent set of proliferative and metabolic stress programs (G2M checkpoint, E2F targets, Mitotic Spindle, mTORC1, Glycolysis, UPR, and ROS; FDR <0.001). This profile highlights cells with high metabolic demand and rapid proliferation occupying geometrically divergent regions of the trajectory, consistent with the NPC/OPC-like cycling progenitor states defined in GBM Neftel et al. (2019).

Applying the identical pipeline to breast cancer preserved this low redundancy (Jaccard overlap = 0.028). Residual TES cells were enriched for EMT-associated transcriptional programs and extracellular matrix organization, consistent with active remodeling states within the tumor microenvironment, whereas residual TDS cells were enriched for distinct lineage-associated transcriptional programs Wu et al. (2021). Although EMT was included as a covariate, residual TES identified additional EMT-related genes. This is expected because the curated signature represents only a subset of the broader transcriptional program; thus, residual enrichment reflects transcriptional programs not fully captured by the canonical covariates rather than implying the complete removal of those biological processes.

Together, despite adjusting for canonical hypoxia, EMT, stemness, and cell-cycle signatures, TES and TDS retained distinct transcriptional associations. TES preferentially marked cells associated with inflammatory remodeling and transcriptional plasticity, consistent with temporal state transitions or inter-compartment boundaries, while TDS was associated with metabolic stress and proliferative activity, consistent with directional instability. This demonstrates that the TES/TDS coordinate system partitions context-specific transitional states—associating with the primary axis of temporal mixing (inflammatory remodeling in GBM; EMT in breast cancer) and fate divergence (proliferative instability in GBM; lineage divergence in breast cancer) specific to the microenvironment. These findings indicate that the two uncertainty metrics capture complementary biological dimensions that are not reducible to conventional cancer-state signatures.

### 3.5 TUF demonstrates parameter stability and sub-sampling robustness

TES and TDS remained robust across a wide range of *k* (10 to 30), with pairwise Spearman correlations ranging from 0.80 to 0.92 (Table 3). Sub-sampling analysis (80% cells, 10 replicates) on pancreatic endocrinogenesis showed high stability in identifying the specific cell types comprising the top 5% uncertain populations (Jaccard *>* 0.95). However, the cluster-level Spearman correlation for TDS (*ρ* = 0.661 *±* 0.232) was lower and more variable than that of TES (*ρ* = 0.829 *±* 0.075). This is a geometrically expected consequence of branch-point cells being inherently rarer than broadly distributed transitional cells; dropping 20% of cells disproportionately impacts the exact local geometry of isolated branch hubs (Table 4).

**Table 3:**
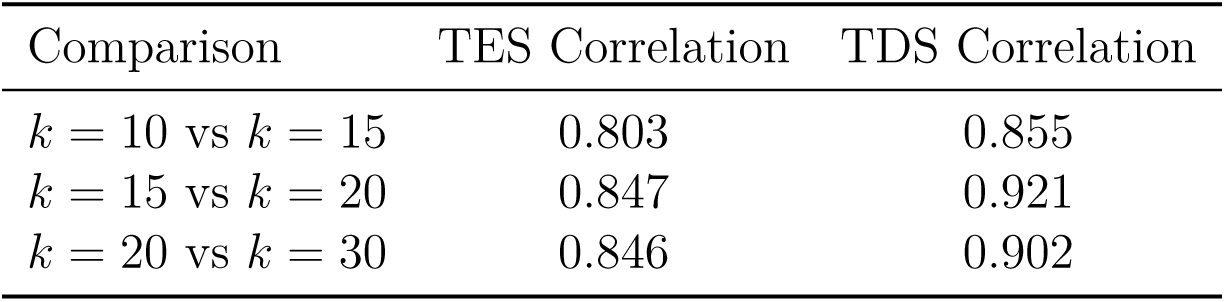
Robustness of TES and TDS to kNN neighborhood size (*k*).

**Table 4:** Subsampling robustness of TUF metrics (80% subsampling, 10 replicates).

| Metric | Cluster-Level Spearman $\rho$ | Top 5% Cell-Type Jaccard |
| --- | --- | --- |
| TES | $0.829 \pm 0.075$ | $0.967 \pm 0.070$ |
| TDS | $0.661 \pm 0.232$ | $0.957 \pm 0.069$ |

Benchmarking confirmed near-linear scaling of TES/TDS computation (Figure 7).

**Figure 7:**
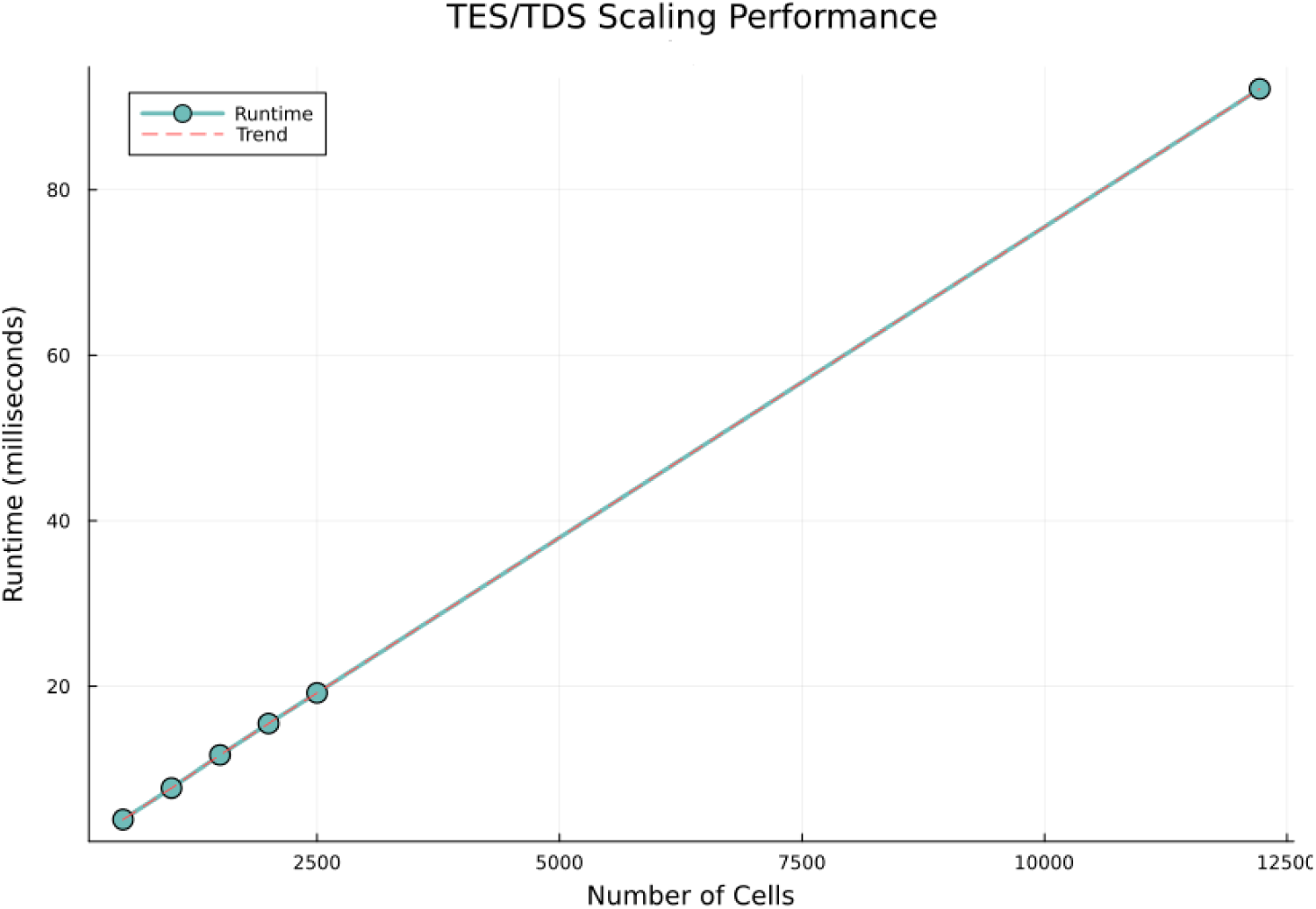
Scaling performance of TES and TDS computation. Runtime (milliseconds) as a function of cell number on the synthetic dataset demonstrates linear scaling.

## 4 Discussion

### 4.1 TUF complements existing trajectory inference methods

TUF provides modular, post-hoc uncertainty quantification that complements existing trajectory inference methods. Conventional algorithms such as DPT Haghverdi et al. (2016), Monocle Trapnell et al. (2014), and Slingshot Street et al. (2018) primarily reconstruct developmental trajectories. TUF quantifies the local uncertainty associated with these inferred trajectories.

Recent methods like Palantir Setty et al. (2019) and CellRank Lange et al. (2022) provide cell-level uncertainty scores, but these are inherently *fate-conditional*: they quantify uncertainty strictly relative to predefined or heuristically inferred terminal states. In contrast, TUF is *fate-agnostic*: it quantifies local geometric and pseudotemporal ambiguity directly from the neighborhood graph. These approaches answer different questions; fate-conditional methods excel at predicting lineage outcomes when terminal states are well-defined, whereas TUF is designed to highlight local geometric incoherence and temporal mixing in contexts where terminal outcomes are ambiguous. TUF acts as a modular, fate-agnostic uncertainty layer that augments, rather than replaces, existing workflows (Table 5).

**Table 5:** Conceptual comparison of TUF with existing trajectory methods.

| Method | Output | Cell score | Local uncert. | Term. states |
| --- | --- | --- | --- | --- |
| DPT Haghverdi et al. (2016) | Pseudotime | ✓ | – | No |
| Monocle Trapnell et al. (2014) | Trajectory | ✓ | – | No |
| Slingshot Street et al. (2018) | Lineage assign. | ✓ | – | No |
| Palantir Setty et al. (2019) | Fate probs. | ✓ | Indirect | Yes |
| CellRank Lange et al. (2022) | Absorption probs. | ✓ | Indirect | Yes |
| **TUF** | **TES + TDS** | ✓ | ✓ | **No** |

### 4.2 Biological interpretation of the TES/TDS coordinate system

The core contribution of TUF is the joint interpretation of TES and TDS as a 2D uncertainty coordinate system. Our cross-tumor analysis reveals that while the low redundancy of these metrics is universal, the biological programs they are associated with are context-dependent. Residual TES is associated with the dominant axis of temporal mixing—inflammatory remodeling and phenotypic transition in glioblastoma, and EMT-associated remodeling in breast cancer. In GBM, this axis seems to be consistent with recently described immune-mediated MES2-like programs Zhang et al. (2025) and CHI3L1–STAT signaling niches Yu et al. (2025). These studies propose that localized immune communication generates transcriptionally heterogeneous microenvironments. The concordance between these observations and TES enrichment suggests that TES captures a fundamental geometric property of the local transcriptomic landscape—the coexistence of heterogeneous transcriptional states during active remodeling—rather than serving as a specific detector of any single molecular pathway. Conversely, residual TDS consistently marks cells exhibiting proliferative and metabolic stress programs (e.g., G2M, mTORC1, Glycolysis), reflecting directional instability in regions of high biosynthetic and geometric divergence characteristic of cycling progenitor states Neftel et al. (2019). More broadly, this suggests that as additional heterogeneous niches (e.g., immune, vascular, or fibroblast-associated) are identified across tissues, TES and TDS may provide general geometry-based metrics for highlighting distinct dimensions of local transcriptional mixing independently of the underlying biological mechanism.

### 4.3 From uncertainty quantification to biological hypothesis generation

Although TUF is designed as a descriptive framework for quantifying local trajectory uncertainty rather than predicting therapeutic response, it provides a principled strategy for prioritizing dynamically heterogeneous cell populations for downstream analysis. Cells exhibiting elevated TES or TDS represent regions of the transcriptomic landscape undergoing active temporal mixing or directional divergence and therefore constitute natural candidates for differential expression, pathway enrichment, gene regulatory network inference, and cell–cell communication analyses.

In cancer and developmental systems, these prioritized populations may help identify cellular states or microenvironmental niches associated with disease progression, tissue remodeling, or lineage diversification. Integration with complementary modalities, including spatial transcriptomics and perturbation-based experiments, may further clarify the functional significance of these high-uncertainty states.

These applications remain prospective and require experimental validation. Rather than providing mechanistic or causal inference, TUF serves as a hypothesis-generating framework that narrows the search space from thousands of cells to biologically meaningful subsets for focused investigation.

### 4.4 Limitations

TUF inherits dependencies from the upstream trajectory inference method: its outputs are only as reliable as the quality of the underlying graph, embedding, and pseudotime. Because TES and TDS quantify geometric properties of inferred trajectories, they should be interpreted as descriptive measures of local uncertainty rather than direct estimates of cell-fate probabilities or developmental potential. In heterogeneous tissues such as tumors, high TUF scores should not be interpreted as evidence of lineage conversion by themselves, as they may represent either within-lineage state transitions or boundaries between distinct cellular compartments. Therefore, interpretation of TUF-highlighted populations requires integration with cell-type annotation and lineage-aware analyses. TDS detects directional incoherence, not exclusively branching events; high TDS can also be triggered by trajectory crossings or boundary artifacts. Therefore, TDS should always be interpreted jointly with TES. Finally, while TUF identifies cells associated with high transcriptional uncertainty, it does not establish causal relationships or predict future cell fates. The framework should be viewed as a hypothesis-generating tool; definitive validation requires experimental lineage tracing or functional perturbations.

### 4.5 Conclusions

By defining uncertainty as a 2D coordinate system rather than a scalar value, TUF enables more nuanced interpretation of cellular decision-making. TUF provides a modular, fate-agnostic uncertainty layer that can be incorporated into diverse single-cell analysis workflows. The development of TUF and its implementation in SiCell.jl illustrate the value of high-performance, open-source tooling in single-cell biology.

## Acknowledgements

The authors acknowledge the use of large language models for spell-checking and rephrasing during the preparation of this manuscript.

## Data and Code Availability

The Trajectory Uncertainty Framework (TUF) is implemented as part of the open-source Julia package **SiCell.jl**, released under the MIT License. The full source code is publicly available on GitHub at https://github.com/Sizerta/SiCell.jl.

SiCell.jl is also registered in the official Julia General Registry and can be installed directly via Julia’s built-in package manager. Comprehensive documentation is available at https://sizerta.github.io/SiCell.jl/. To ensure full reproducibility of the cross-pipeline validation results, Python/Scanpy scripts are provided within the repository. All datasets are publicly available (Table 1).

## Author Contributions

M.M. conceived the study, developed the SiCell.jl software and the TUF mathematical framework, performed the computational analyses, and wrote the manuscript.

Z.M. contributed to the biological interpretation of the results, helped design and refine the biological analyses, assisted in the interpretation of pathway enrichment, and reviewed and edited the manuscript.

T.I. contributed to the biological interpretation of the findings, particularly from a cell-state and systems biology perspective, provided input on future research directions, and reviewed and edited the manuscript.

All authors read and approved the final manuscript.

