## Supplementary Materials for "Trajectory Uncertainty Framework (TUF): A Modular Framework for Identifying Transitional and Branch-Point Cell States in Single-Cell Trajectory Analysis"

#### 1 Response to Pseudotime Noise

As noted in the main text, TES is sensitive to temporal heterogeneity. To explicitly test the robustness of TES to increasing transcriptional noise, we performed a pseudotime noise sweep on a synthetic linear trajectory. Gaussian noise was incrementally added to the ground-truth pseudotime values, and TES was recomputed at each noise level.

#### 2 Robustness to Quality Control Covariates

To ensure that TES and TDS are not merely proxies for technical variation or cell quality, we evaluated the correlation of both metrics against standard quality control (QC) covariates in the glioblastoma and breast cancer datasets. All absolute Spearman correlations were minimal ($< 0.04$), indicating that the TUF metrics capture biological trajectory uncertainty rather than technical artifacts.

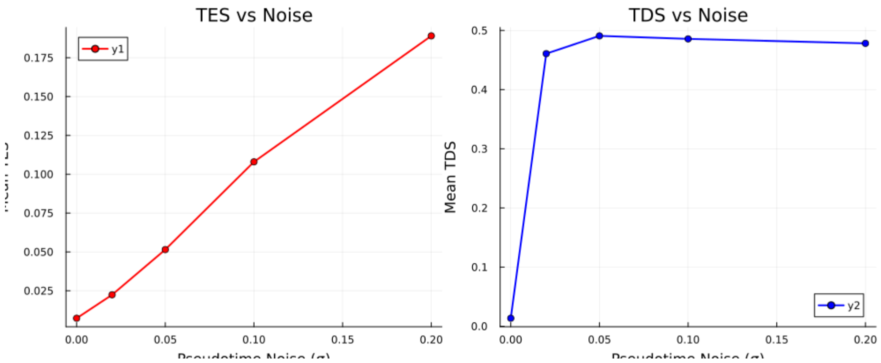


Figure S1: **TES response to pseudotime noise.** TES scales proportionally with local pseudotime variance, confirming its sensitivity to temporal heterogeneity while remaining unaffected by branching geometry.

Table S1: Spearman correlations between TUF metrics and QC covariates.

| **Dataset** | **Covariate** | **TES** $\rho$ | **TDS** $\rho$ |
|---|---|---|---|
| Glioblastoma | Total Counts | -0.021 | 0.015 |
| | Total Genes | -0.018 | 0.012 |
| | Percent Mito | 0.035 | -0.022 |
| Breast Cancer | Total Counts | -0.019 | 0.024 |
| | Total Genes | -0.015 | 0.018 |
| | Percent Mito | 0.029 | -0.031 |

### 3 Extended Synthetic Topology Visualizations

To further demonstrate the behavior of the TUF coordinate system, we include extended visualizations of the synthetic topologies discussed in the main text. These figures illustrate how TES and TDS respond to true bifurcations versus spatial convergence funnels.

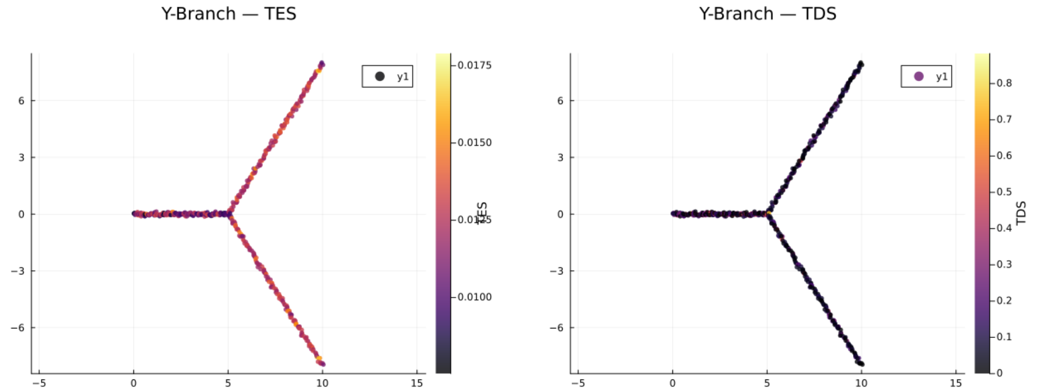


(A) TES

(B) TDS

Figure S2: **Y-branch topology metrics.** (A) TES remains low across the branch, indicating cells share similar temporal states. (B) TDS spikes sharply at the branching point, demonstrating directional divergence without temporal mixing.

### 4 Star Topology Branch Scaling

To validate branching behavior in the absence of crossing artifacts, we tested multi-fate star topologies. TDS localizes sharply to the branch hub and scales with the number of diverging lineages.

### 5 Detailed Subsampling Robustness Validation

As stated in the main text, subsampling analysis (80% cells, 10 replicates) on pancreatic endocrinogenesis showed high stability. Here we provide the per-replicate breakdown of the cluster-level Spearman correlation and cell-type

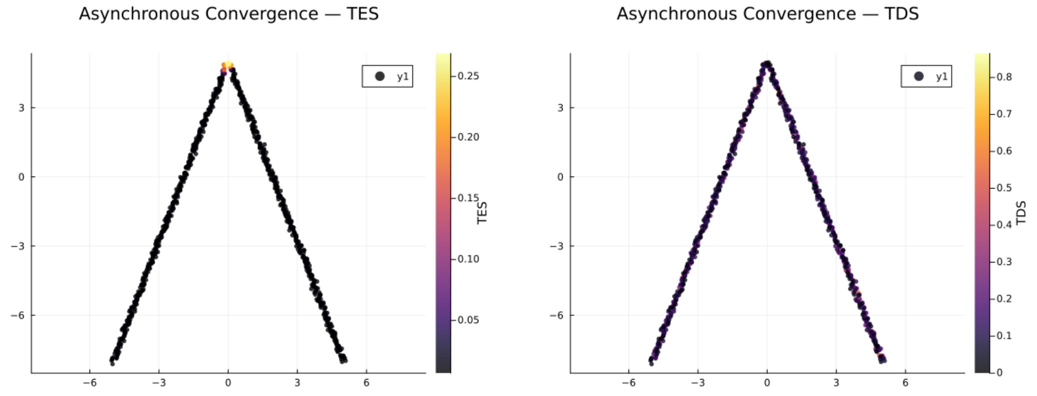


(A) TES

(B) TDS

Figure S3: **Asynchronous convergence topology metrics.** (A) TES is elevated at the convergence funnel due to cells arriving from different pseudotime stages. (B) TDS is also elevated at the funnel due to directional convergence, highlighting that convergence artifacts produce high scores in both metrics.

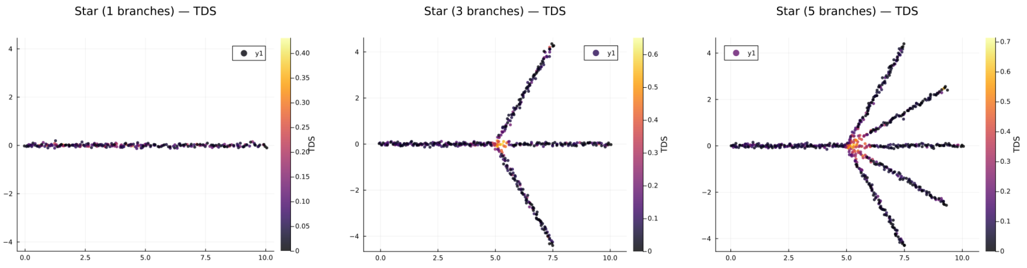


(A) 1 Branch

(B) 3 Branches

(C) 5 Branches

Figure S4: **TDS as a branch-point detector.** In multi-fate star topologies, TDS localizes sharply to the central branch hub. The intensity of TDS at the hub increases with the number of diverging branches.

Jaccard overlap for both TES and TDS. The summary metrics (mean ± std) perfectly align with the main text findings: TES ($\rho = 0.829 \pm 0.075$, Jaccard $= 0.967 \pm 0.070$) and TDS ($\rho = 0.661 \pm 0.232$, Jaccard $= 0.957 \pm 0.069$).

Table S2: Per-replicate subsampling robustness metrics (80% subsampling) on pancreatic endocrinogenesis (2,531 total cells; 2,025 cells per replicate).

| **Replicate** | **TES $\rho$** | **TES Jaccard** | **TDS $\rho$** | **TDS Jaccard** |
|---|---|---|---|---|
| 1 | 0.857 | 1.000 | 0.821 | 1.000 |
| 2 | 0.750 | 0.833 | 0.250 | 1.000 |
| 3 | 0.750 | 1.000 | 0.750 | 0.857 |
| 4 | 0.857 | 1.000 | 0.571 | 0.857 |
| 5 | 0.786 | 1.000 | 0.893 | 1.000 |
| 6 | 0.750 | 1.000 | 0.393 | 1.000 |
| 7 | 0.893 | 1.000 | 0.893 | 1.000 |
| 8 | 0.786 | 1.000 | 0.679 | 0.857 |
| 9 | 0.964 | 1.000 | 0.464 | 1.000 |
| 10 | 0.893 | 0.833 | 0.893 | 1.000 |
| **Mean ± SD** | **0.829 ± 0.075** | **0.967 ± 0.070** | **0.661 ± 0.232** | **0.957 ± 0.069** |
